# Generation of human induced pluripotent stem cell (hiPSC) lines from patients with extreme high and low polygenic scores for QT interval

**DOI:** 10.1101/2023.11.03.565486

**Authors:** Devyn Mitchell, Rizwan Ullah, Loren Vanags, Alex Shen, Luke Jones, Matthew O’Neill, Giovanni Davogustto, Christian Shaffer, Dan Roden, Ben Shoemaker, Hollie Williams, Teresa Strickland, Taylor Agee, Christopher Johnson, Brett Kroncke

## Abstract

Long QT syndrome (LQTS) is an inherited cardiac arrhythmia syndrome with congenital and drug-induced presentations and known monogenic and polygenic contributions. LQTS represents a significant clinical challenge due to its complex genetic underpinning and propensity for fatal arrhythmias. In this study, we generated induced pluripotent stem cells (iPSCs) reprogrammed from peripheral blood mononuclear cells (PBMCs) of six patients with extreme polygenic scores for short and long corrected QT intervals. iPSC lines were rigorously validated for genomic integrity through karyotyping and targeted mutation analysis specific to a lengthened or shortened QT interval. Pluripotency was confirmed by expression of key markers TRA 1-60, TRA 1-81, SOX2, OCT4, NANOG, and REX1 via quantitative PCR and immunofluorescence. Subsequent cardiac induction successfully generated cardiomyocytes that were further characterized. This patient-specific approach will enable us to better understand variable expressivity and penetrance of LQTS. Rigorously validated iPSC lines serve as a vital resource for elucidating the molecular mechanisms underlying LQTS. Our study provides a robust and clinically relevant resource to facilitate our understanding the genetic and cellular complexity of LQTS.

## Introduction

Long QT syndrome (LQTS; OMIM: 613688) is associated with sudden unexplained death in young populations. LQTS is an inherited cardiac arrhythmia with congenital and drug-induced presentations defined by a prolongation of the corrected QT interval on an electrocardiogram.^1, 2^ The QT interval, corrected for heart rate (QTc), is 30-40% heritable,^3, 4^ with narrow-sense, autosomal common single nucleotide polymorphisms (SNP) contribution estimated around 20%;^5, 6^ the SNP contribution to LQTS is similar, around 15%.^7^ Until recently, LQTS had been thought of as largely a monogenic disorder with approximately 75% of congenital LQTS probands carrying a variant in one of three cardiac ion channel genes: *KCNQ1*, *KCNH2*, and *SCN5A*.^8, 9, 10, 11, 12^ These three gene products help control the duration and profile of the cardiomyocyte action potential, mechanistically linking all three to LQTS.^13, 14^ Since these initial associations were uncovered, 14 additional genes have been proposed as monogenic causes of LQTS ^15^. However, only three are considered *Definitive* in Clinical Genome Resource (*CALM*1/2/3).

We identified six different patients (Table 4) with a diverse set of SNPs (single nucleotide polymorphism) associated with varying degrees of polygenic propensity for a high, medium, or low QT interval. The polygenic score (PGS) was derived from 68 independent SNPs at the 35 loci which collectively explain around 10% of variance in QT interval ^24^. The six patient lines’ PGS percentile were calculated using the standard deviation from these 68 SNPs from each patient line from a previously assembled registry comprising 2,899 genotyped participants ^25, 26^. The percentile PGS values for each line are two from low (0.01% PGS, 0.33% PGS), two from middle (50.2% PGS, 63.5% PGS), and two from high (99.4% PGS, 99.7% PGS). All six patient lines were reprogrammed from patient peripheral blood mononuclear cells (PBMCs) into induced pluripotent stem cells (iPSC). Stem cells are undifferentiated cells with the ability to self-renew and differentiate into a variety of cell types, making them attractive candidates for cell-based therapies and in vitro studies of human pseudo-tissue. However, traditional methods for generating stem cells, such as through embryonic or adult tissue sources, are often limited in quantity and quality. Therefore, the breakthrough in generating iPSCs through somatic cell reprogramming has ushered in a new era for the study of cardiomyocytes. This innovative approach enables the creation of patient-specific cardiomyocytes derived directly from iPSCs, which are reprogrammed from patient blood. The process of reprogramming somatic cells into iPSCs involves the introduction of a defined set of transcription factors, typically Oct4, Sox2, Klf4, Nanog, Lin28 and c-Myc, into the cells ^17, 18^. These factors can induce the expression of genes associated with pluripotency, leading to the formation of iPSCs.

iPSCs can be differentiated into a growing number of cell types, including atrial and ventricular cardiomyocytes. Cardiomyocytes derived from iPSCs offer distinct advantages over other cell sources for cardiac studies due to their close resemblance to native human cardiomyocytes, including similar transcriptional and protein expression profiles. Differentiation of iPSCs into cardiomyocytes involves the recapitulation of the cardiac developmental program, which is regulated by the coordinated expression of various transcription factors, signaling molecules, and epigenetic modifications ^19^. The differentiation process is typically initiated by treating iPSCs with small molecules or growth factors that modulate key signaling pathways involved in cardiac development, such as the Wnt, BMP, and FGF pathways ^20^. Subsequent manipulation of culture conditions, such as changes in medium composition and substrate stiffness, can further enhance the efficiency and purity of cardiomyocyte differentiation ^21, 22^. Other protocols involve the use of transcription factors or direct reprogramming to bypass the pluripotent state and directly induce cardiac differentiation ^23^.

In the present study, we have successfully reprogrammed cell lines obtained from patient samples, an advancement that enables us to faithfully replicate cellular characteristics and disease-associated phenotypes, which are otherwise challenging to investigate directly in patients. Our main objective was to establish iPSC lines as a platform probing them mechanisms of common genetic variation on QT interval presentation and its intricate interplay with rare, large-effect variants. Through careful optimization and meticulous characterization, we established six iPSC lines that displayed key characteristics of iPSC-CMs. All six iPSC lines generated displayed normal embryoid stem cell morphology and exhibited normal karyotyping after reprogramming. iPSC lines underwent STR (short tandem repeated DNA sequences) analysis with seven commonly used loci D151656, D165539, D75820, D135317, THO1, TPOX, and D55818 to validate each line was derived from patient PBMCs. Mutation analysis of ten different loci ensure the SNP of each iPSC line were consistent with previous sequencing after reprogramming. Each iPSC line displayed the expression of pluripotent markers REX1, NANOG, OCT4, SOX2, and house-keeping gene GAPDH through PCR analysis. Immunofluorescence staining of all iPSC lines showed expression of pluripotent markers TRA1-60, OCT4, TRA1-81, and NANOG. Additionally, immunofluorescence staining of iPSC-CMs demonstrated expression of α-Actinin and MLC-2V.

## Methods and Materials

### Peripheral Blood Mononuclear Cells (PBMC) isolation from blood samples

Blood samples were collected from patients to generate induced pluripotent stem cells (iPSCs). The samples were spun in the centrifuge (Allegra X-14R), at 2,000 g for 15 minutes then gradually slowed to a stop as to not disturb the buffy layer. Extreme care was taken before opening the tubes with careful sterile technique.

The plasma was separated from the buffy coat and red blood cells. Typically, two 5 mL vials of blood were collected from each patient. The plasma was then carefully transferred into three cryo tubes using transfer pipettes, being careful not to disturb the buffy coat.

The buffy coat, which contains peripheral blood mononuclear cells (PBMCs), was processed by transferring it into a 15 mL conical tube and adding enough Dulbecco’s phosphate-buffered saline (DPBS) (Cat# 14190-144 Thermofisher Scientific) to make a total volume of 14 mL. The tube was then spun at 300 g for 10 minutes. Supernatant was aspirated off, and cell pellet was resuspended in 5 mL of DPBS. One Countess cell counting chamber slide (Cat#C10283 Invitrogen) per patient was prepared for counting cells and placed under the hood. The cells were counted twice for each slide by Countess 3 Automated Cell Counter (Cat#AMQAX2000 Invitrogen), and the number of cells and percentage of live cells were recorded. 3 million cells were stored in each tube.

To prepare the cells for cryo preservation, the cell suspension was spun at 300 g for 5 minutes. While the cells were spinning, the freezing media was prepared by adding 10% DMSO to Fetal Bovine Serum (FBS) (Cat#A31606-01 Gibco), 1 mL of freezing media per 3-4 million cells. The DPBS was aspirated as close to the pellet as possible, and the freezing media was added to the pellet. The pellet was gently resuspended in the freezing media, and 1 mL of the cell suspension was aliquoted into each labeled Nalgene Cryoware Cryogenic Vials (cat#5000-0020 Thermofisher). The tubes were then placed in a cylindrical container with 70% isopropanol. The container was kept in a - 80°C freezer for at least 24 hours for slow freezing, then PBMC samples were moved into liquid nitrogen dewar for long term storage.

### Generation of induced pluripotent stem cells (iPSCs) from peripheral blood mononuclear cells (PBMCs)

To generate iPSCs from peripheral blood mononuclear cells (PBMCs) the cells were thawed in a 37°C water bath until fully thawed. For this labor intensive process, we began this protocol on Monday, Wednesday, or preferably Friday. PBMCs were then added to 9 mL of SFM media (Table 1) and spun at 300 g for 5 minutes. Once spun down, the supernatant was aspirated as close to cell pellet as possible. The cell pellet was then resuspended in 1 mL of MNC media (Table 1) at a concentration of 2.5 million to 4 million cells per mL, leading to the erythroblast enrichment and maintenance. The MNC media was made fresh daily, and all reagents were stored at their respective temperatures until use. Dexamethasone was kept as a 12 mM stock and diluted 1:3 with water for a working concentration of 4 mM. Recombinant Human Erythropoietin (EPO) and Human Holo Transferin (HHT). It was noted that all reagents were only used twice due to freeze/thaw degradation. After adding all necessary reagents, the MNC media was filtered using a 20 µm syringe filter (Cat#431229 Coring). Erythroblast enrichment and maintenance were performed starting the following Monday of thawing. Live cell counts were taken every other day using trypan blue stain 0.4% (Cat#T10282 Invitrogen), and a reduction was noted in cell count in the first few days. Media changes were performed every other day, following an M-W-F schedule. Cells were spun at 300 g for 5 minutes and plated in MNC media at a density of 1 X 10^6^ cells per mL. Erythroblasts were ready for reprogramming once cells became reddish-brown with at least 2 million cells which typically occurred within 10-14 days of initial plating.

**Table 1.**
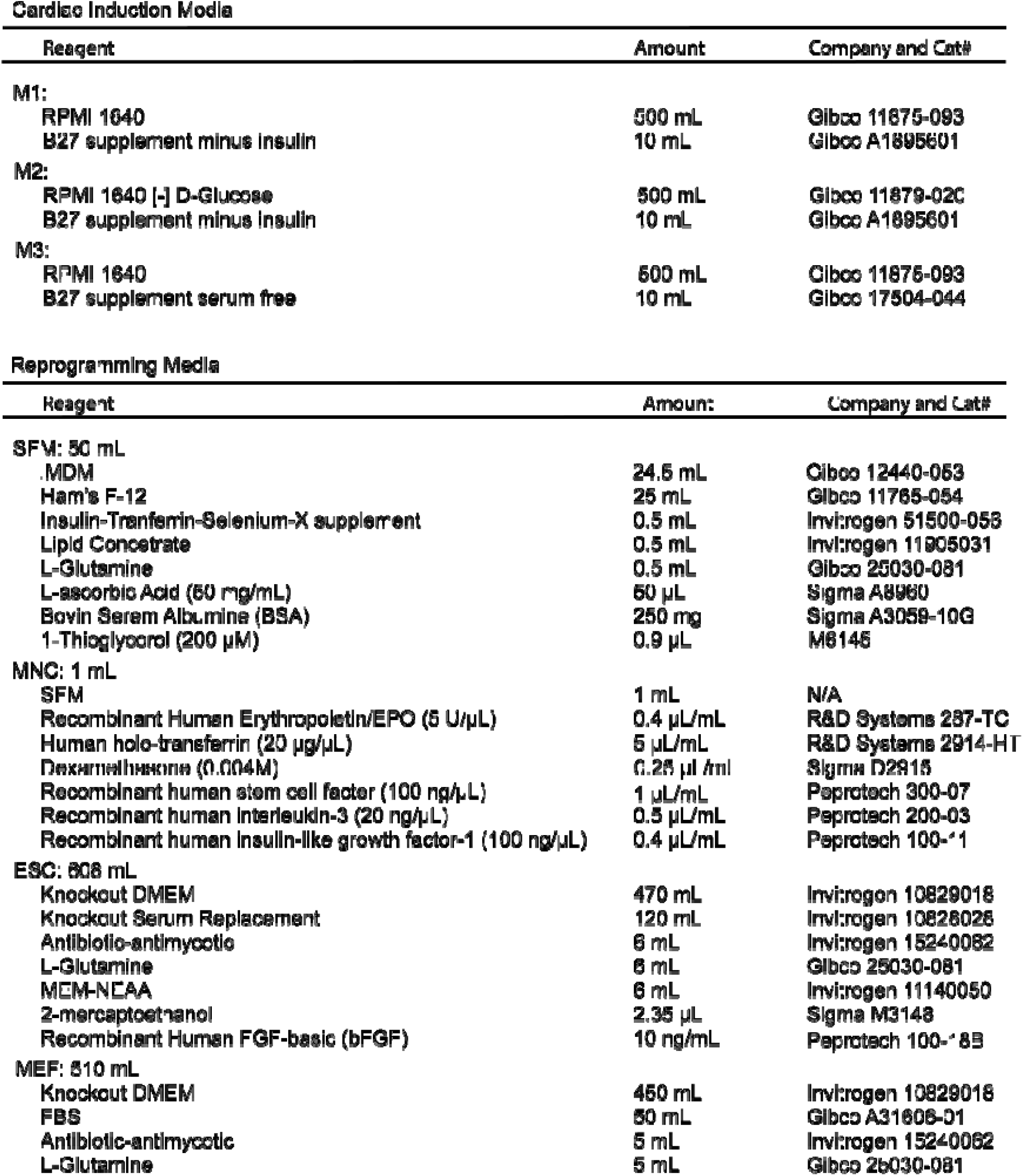

On Day 0 which was set on Monday, 2 million live cells were transferred to a 15 mL conical tube with 10 mL of Dulbecco’s phosphate-buffered saline (DPBS) (Cat# 14190- 144 Thermofisher Scientific) and centrifuged at 300 g for 5 min. The supernatant was aspirated, and the pellet was resuspended in 2 mL DPBS before transferring equal parts to 2 eppendorff tubes (1 million live cells each). These tubes were centrifuged at 300 g for 5 min, and the DPBS was aspirated before resuspending the pellet in the reprogramming cocktail from Epi5 iPSC Reprogramming Kit (Cat#A15960 Invitrogen) along with Neon Transfection System 10 μL Kit (Cat#MPK1096 Invitrogen). The reprogramming cocktail was prepared by adding 10 μL buffer T, 1 μL each of the reprogramming vectors, and 1 μL of EBNA and dominant negative p53, which were kept on ice. The cell pellet was resuspended in 12 μL reprogramming cocktail. NEON electroporation parameters were 1650 V, 10 ms, and 3 pulses, and the transfected cells were deposited into a well of a 12-well plate containing 1 mL fresh MNC media.

Day 1, gelatin plates were prepared and irradiated CF1 Mouse Embryonic Fibroblasts, (MEFs) (Cat# A34181 Gibco), were thawed and plated. Gelatin plates were prepared by heating stock gelatin (Cat# G1393 Sigma) in a 37% water bath, mixing it with DPBS without Ca and Mg (0.1% gelatin/DPBS) in a 50 mL conical tube, and adding 2 mL of gelatin to each well of a labeled 6-well plate. The plate was covered and allowed to sit for at least 20 minutes for the gelatin to settle to the bottom of the well. After aspirating the gelatin, the plate was left uncovered in the back of the hood. MEFs, which had been cryopreserved in MEF media (Table 1) with 10% DMSO, were thawed in a 37°C bath until no longer frozen and then transferred to 9-10 mL of MEF media for a wash. The cells were spun at 300 g for 5 minutes at room temperature and resuspended in 2 mL of MEF media to achieve a concentration of 333,000 to 500,000 irradiated MEF cells per well. The cells were evenly distributed in the well using a back-and-forth and side-to-side motion and were checked under a microscope to ensure even distribution.

Day 2, the cells that underwent reprogramming were transferred into a 15 mL conical containing 9 mL of DPBS. Cells were then spun down at 300 g for 5 minutes and resuspended in a 6-well plate containing irradiated MEF cells and 1 mL of MEF media per well. The entire plate was spun (Allegra X-14R) at 200 g for 30 min until it gradually came to a stop, which helped the erythrocytes settle onto the MEFs.

On Day 3, 1 mL MNC media was added to each well, and on Day 4 and Day 5, 1 mL ESC media (Table 1) was added per well without removing any media.

From Day 7 to Day 13, complete media changes were carried out with 2 mL ESC media per well, and on Day 15, the media was switched to mTeSR Plus media (Cat#100-0276 StemCell Technologies) with 2 mL per well. Finally, until hiPSC colonies develop, complete media changes were carried out with mTeSR, and colonies were picked into a Cultrex stem cell qualified reduced growth factor basement membrane extract (cat# 3434-005-02 R&D systems Inc) coated 24-well plate for expansion. Unpurified colonies were frozen down for safety, and colonies were picked and passaged until hiPSC lines were as pure as possible before karyotyping.

### Karyotyping

To validate chromosomal integrity from reprogramming, generated human induced pluripotent stem cells (hiPSCs) were submitted to Genetics Associates for karyotyping analysis. hiPSCs were plated onto Cultrex stem cell qualified reduced growth factor basement membrane extract (cat# 3434-005-02 R&D systems Inc) coated 25 mL flask (cat#353018 Falcon) and submitted for pick-up by Genetic Associates. Colonies were selected after ‘normal’ karyotyping. (Figure 4A for chromosome image, full set of images in supplementary data)

### Mutation Analysis

Approximately 1 million human induced pluripotent stem cells (hiPSCs) were collected and spun in the centrifuge at 6.8 g for 5 minutes. The supernatant was aspirated from the cell pellet, and then resuspended with 100 µL per 1 million cells of QuickExtract (Pt#SS000035-D1 Biosearch Technologies). The hiPSCs were then run in a thermocycler for three consecutive cycles. The first cycle was at 65 °C for 15 minutes, the second cycle was at 98°C for 10 minutes, and the third cycle was at 4°C as a holding phase. The resulting genomic DNA extract from the hiPSCs was amplified using a Polymerase Chain Reaction (PCR) protocol by preparing each PCR tube, 12.5 µL of Promega Master Mix (Cat# M7502 Promega), 6.5 µL of Molecular Biology Grade water (ref#46-000-CI Corning), 2 µL of 10 mM forward primer (Table 3), 2 µL of 10 mM reverse primer (Table 3), 2 µL of DNA from QuickExtract (See Table 3 for primer list) were added to make a total volume of 25 µL in each tube. Samples then underwent the following PCR protocol: 98°C for 40 seconds, followed by 10 cycles of 98°C for 10 seconds, 65°C-55°C for 30 seconds, and 72°C for 1 minute, followed by 22 cycles of 98°C for 10 seconds, 60°C for 30 seconds, 72°C for 1 minute, and finally 1 cycle of 72°C for 5 minutes, and 4°C as a holding phase. 5 μL of PCR product was then run on a 1% agarose gel at 130 V for 30 minutes. The remaining tubes were enzymatically purified using ExoSAP reagent (#1338674 thermofisher). 1 µL of the ExoSAP enzyme was added per 10 µL of PCR product, and the PCR tubes were put in the thermocycler to run at 37°C for 30 minutes and then 80°C for 15 minutes. 10 µL of each purified sample was submitted for Sanger sequencing (Genewiz; Figure 4B, full set of mutation analysis images in supplementary data)

### STR Analysis

Short-tandem repeats (STR) analysis was used to compare patient PBMCs to PBMC-derived iPSCs for validation. Approximately 1 million hiPSCs and PBMCs from each patient line were collected and spun in the centrifuge at 6800 g for 5 minutes. The supernatant was aspirated from the cell pellet. The cell pellet was then resuspended with 100 µL per 1 million cells of QuickExtract (Pt#SS000035-D1 Biosearch Technologies). The hiPSCs and PBMCs were then run in a thermocycler for one round of three consecutive cycles. The first cycle was at 65 °C for 15 minutes, the second cycle was at 98°C for 10 minutes, and the third cycle was at 4°C as a holding phase.

Once the Genomic DNA Extract was removed from the thermocycler, the genomic DNA of the hiPSCs and PBMCs was amplified using a Polymerase Chain Reaction (PCR) protocol. To prepare each PCR tube, 12.5 µL of Promega Master Mix (Cat# M7502 Promega), 6.5 µL of Molecular Biology Grade water (ref#46-000-CI Corning), 2 µL of forward primer 10 mM (Table 3), 2 µL of reverse primer 10 mM (Table 3), 2 µL of DNA from QuickExtract were added to make a total volume of 25 µL in each tube. Each tube was then placed in the thermocycler and the thermocycler was run at 98°C for 40 seconds, followed by 10 cycles of 98°C for 10 seconds, 65°C-55°C for 30 seconds, and 72°C for 1 minute, followed by 22 cycles of 98°C for 10 seconds, 60°C for 30 seconds, 72°C for 1 minute, and finally 1 cycle of 72°C for 5 minutes, and 4°C as a holding phase. PCR products were then run on a 3% agarose gel at 130 V for 1 hour then gel was analyzed. (Figure 4D for Gel image, full set of images in supplementary data)

**Table 2.**
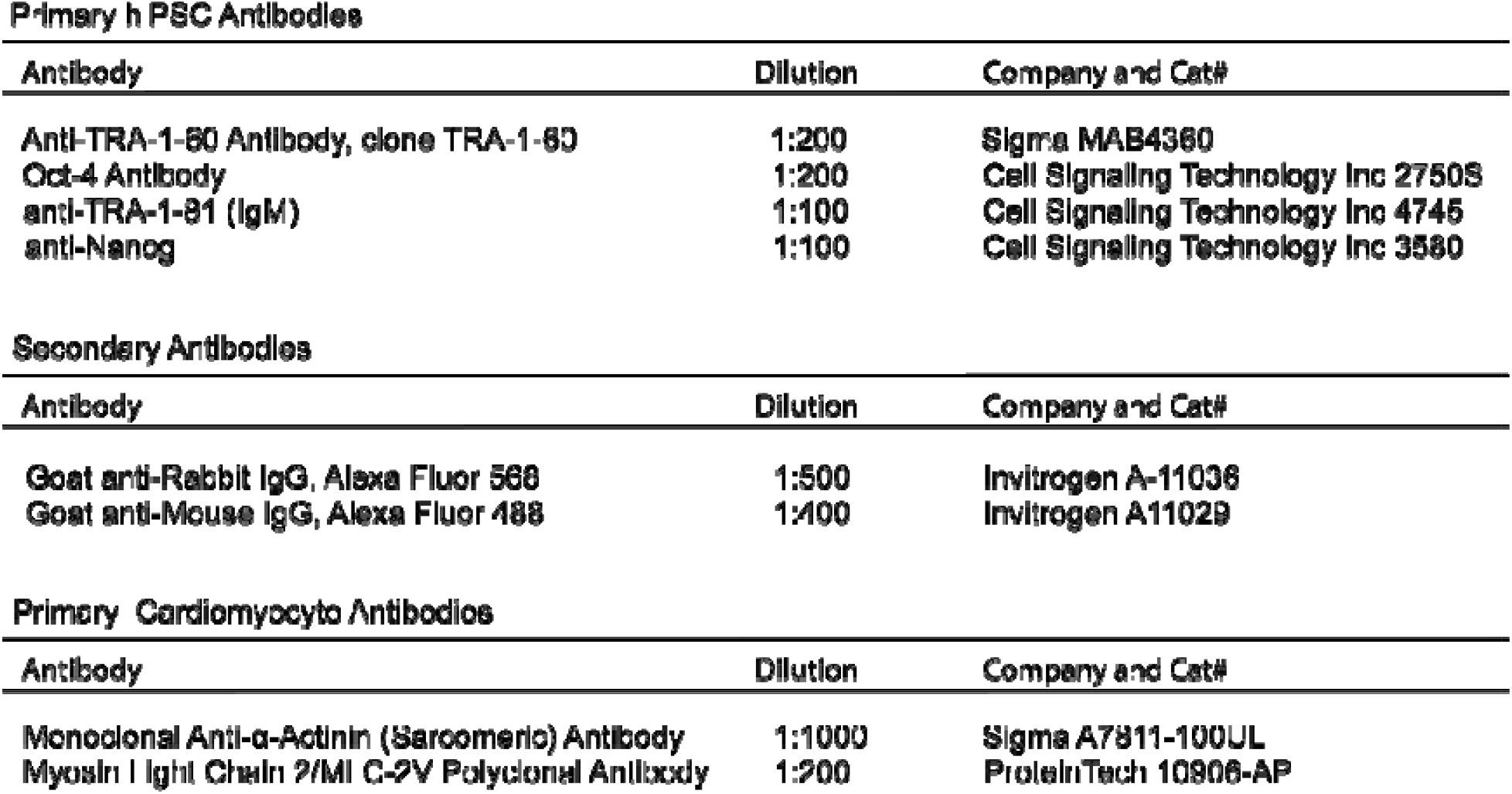

**Table 3.**
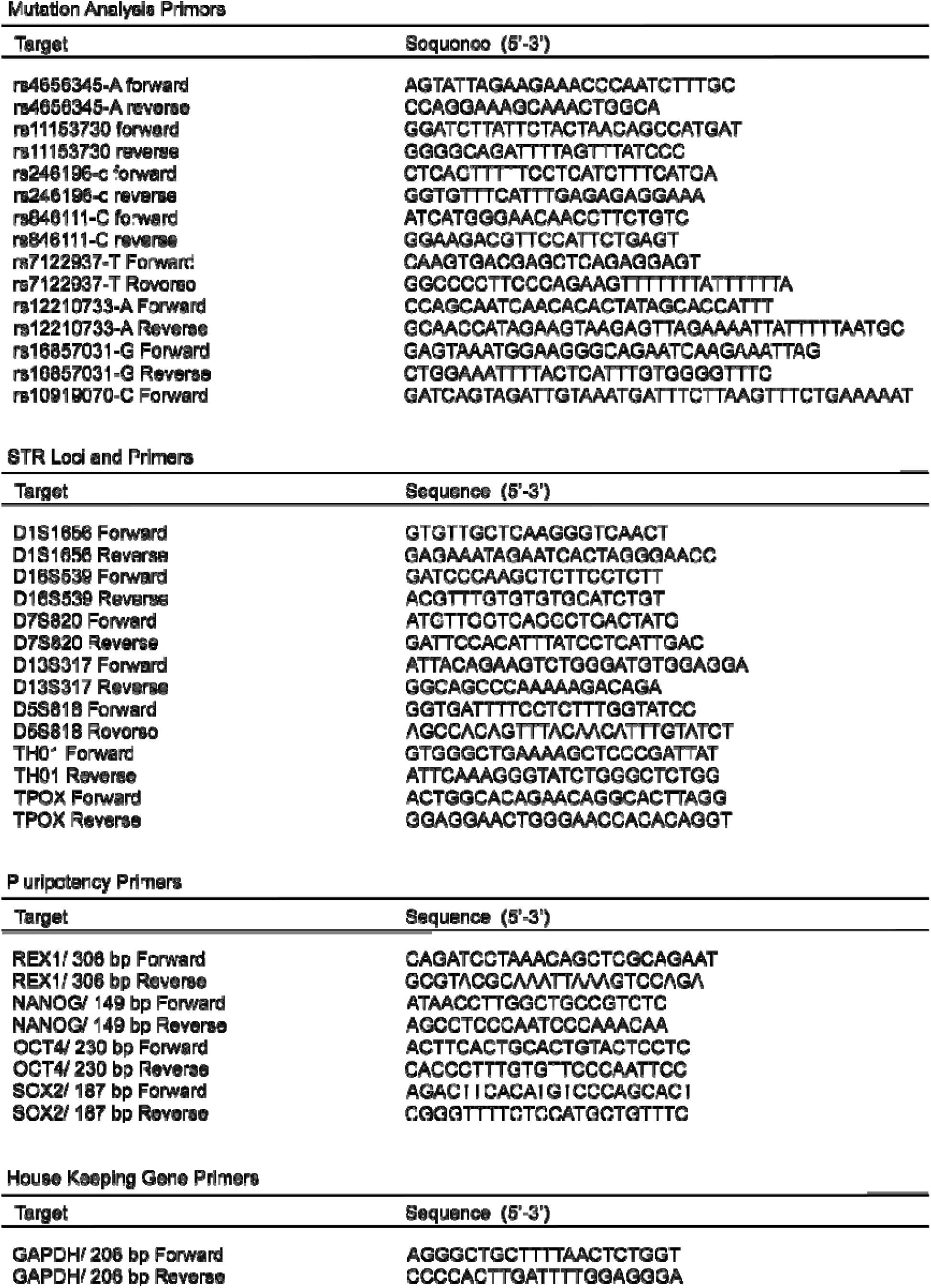

**Table 4.** IPSC Line Information

| QT PGS | Sex | Age | Ethnicity | Cardiac Phenotypes |
| --- | --- | --- | --- | --- |
| 0.01% | Male | 91 | White/Non-hispanic | AFib, DiTdP |
| 0.33% | Male | 77 | Non-hispanic | AFib, Control for DiTdP on Dofetilide |
| 50.2% | Female | 71 | White/Unknkown | AFib, DiTdP |
| 63.5% | Female | 90 | White/Black/Non-hispanic | AFib |
| 99.4% | Male | 70 | White/Non-hispanic | AFib, IgM kappa MGUS |
| 99.7% | Male | 66 | White/Non-hispanic | AFib, HCM, MVD, NICM |

AFib - Atrial fibrillation
DiTdP - drug Induced Torsades de Pointes
HCM - Hypertrophic cardiomyopathy
IgM kappa MGUS - Immunoglobulin M kappa monoclonal gammopathy of undetermined significance
MVD - Coronary microvascular disease
NICM - Nonischemic cardiomyopathy

### Pluripotency Gene Expression

Approximately 1 million hiPSCs were collected and spun in the centrifuge at 6800 g for 5 minutes. The supernatant was aspirated from the cell pellet. The RNeasy Mini Kit from QIAGEN (Cat.#74104) was used to extract RNA from between 1 million and 5 million hiPSCs. In order to convert the extracted RNA from those cells into cDNA for amplification, we used the SuperScript III First-Strand Synthesis System for RT-PCR (Cat.# 18080-051). The cDNA was then amplified using a Polymerase Chain Reaction (PCR) protocol. To prepare each PCR tube, 12.5 µL of Promega Master Mix (Cat# M7502 Promega), 6.5 µL of Molecular Biology Grade water (ref#46-000-CI Corning), 2 µL of forward primer 10 mM (Table 3), 2 µL of reverse primer 10 mM (Table 3), 2 µL of cDNA product were added to make a total volume of 25 µL in each tube. Each tube was then placed in the thermocycler and the thermocycler was run at 98°C for 40 seconds, followed by 10 cycles of 98°C for 10 seconds, 65°C-55°C for 30 seconds, and 72°C for 1 minute, followed by 22 cycles of 98°C for 10 seconds, 60°C for 30 seconds, 72°C for 1 minute, and finally 1 cycle of 72°C for 5 minutes, and 4°C as a holding phase. PCR samples were then run on a 1% agarose gel at 130 V for 1 hour and the gel was subsequently analyzed. (Figure 4C for Gel images)

### hiPSC Confocal Microscopy

Generated hiPSC lines were layered onto a cultrex stem cell qualified reduced growth factor basement membrane extract (cat# 3434-005-02 R&D systems Inc) coated 6-well plate with cover slips (cat#12-542-BP FisherBrand) in each well. For each 6-well cells, we added 2 mL of 4% PFA (cat#28906 Thermofisher) in PBS (cat#10010-023 Gibco) for 30 minutes at room temperature (RT). After fixation, cells were washed three times with PBS to remove residual PFA. Next, cells were permeabilized with 0.2% Triton X-100 (cat#T8787 Sigma) in PBS for 20 minutes at RT. Cells are then blocked with 2 mL blocking solution made of 5% BSA (cat#A3059-10G Sigma) and 0.05% Triton X-100 in PBS for 2 hours at RT or 4°C on a shaker.

Primary antibodies against stem cell markers (TRA1-60 (Table 2), OCT4 (Table 2), NANOG (Table 2), TRA1-81 (Table 2)) were added with 500 μL of blocking solution to the cells and incubated overnight at 4°C on a shaker. After incubation with primary antibodies, cells are washed three times with 0.05% Triton X-100/PBS for 5 minutes each. Secondary antibodies, Goat anti-Mouse IgG (H+L) Highly Cross-Absorbed Secondary Antibody Alexa Fluor 488 (Table 2) and, Goat anti-Rabbit IgG (H+L) Highly Cross-Absorbed Secondary Antibody Alexa Fluor 568 (Table 2) were then added with 500 μL of blocking solution and incubated for 4 hours at RT or overnight at 4°C on a shaker. After secondary antibody incubation, cells were washed three times with 0.05% Triton X-100/PBS for 5 minutes each. Next, 1 mL of Hoechst dye 2 ug/mL (cat#B1155 Sigma) in PBS was added to cells for nuclear counterstaining and incubated for 5 minutes at RT. After Hoechst dye incubation, cells were washed three times with PBS for 5 minutes each. Cells were then placed onto a glass slide (cat#12-550-433 Fisherbrand) with aqua hold 2 pen (Cat# 9804-02 Scientific Device) to seal slide, and then a cover slip was added with 50% glycerol (cat#17904 Sigma) in PBS. The edges of the cover slip were sealed with nail polish, and the slide was allowed to dry overnight at 4°C. Stained hiPSC slides were then imaged on Zeiss LSM 710 confocal microscope (Figure 2).

**Figure 1.**
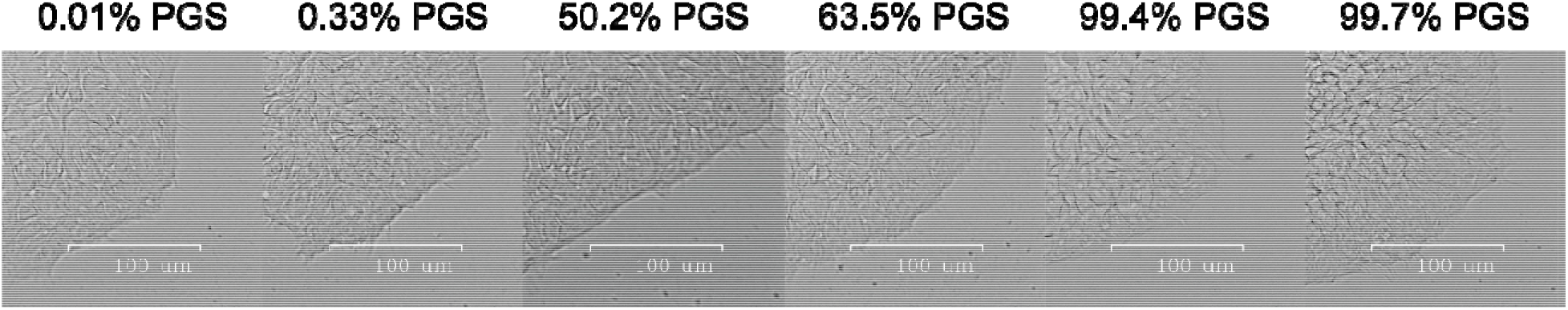
All six iPSC generated lines showed normal Embryonic Stem-like morphology.

**Figure 2.**
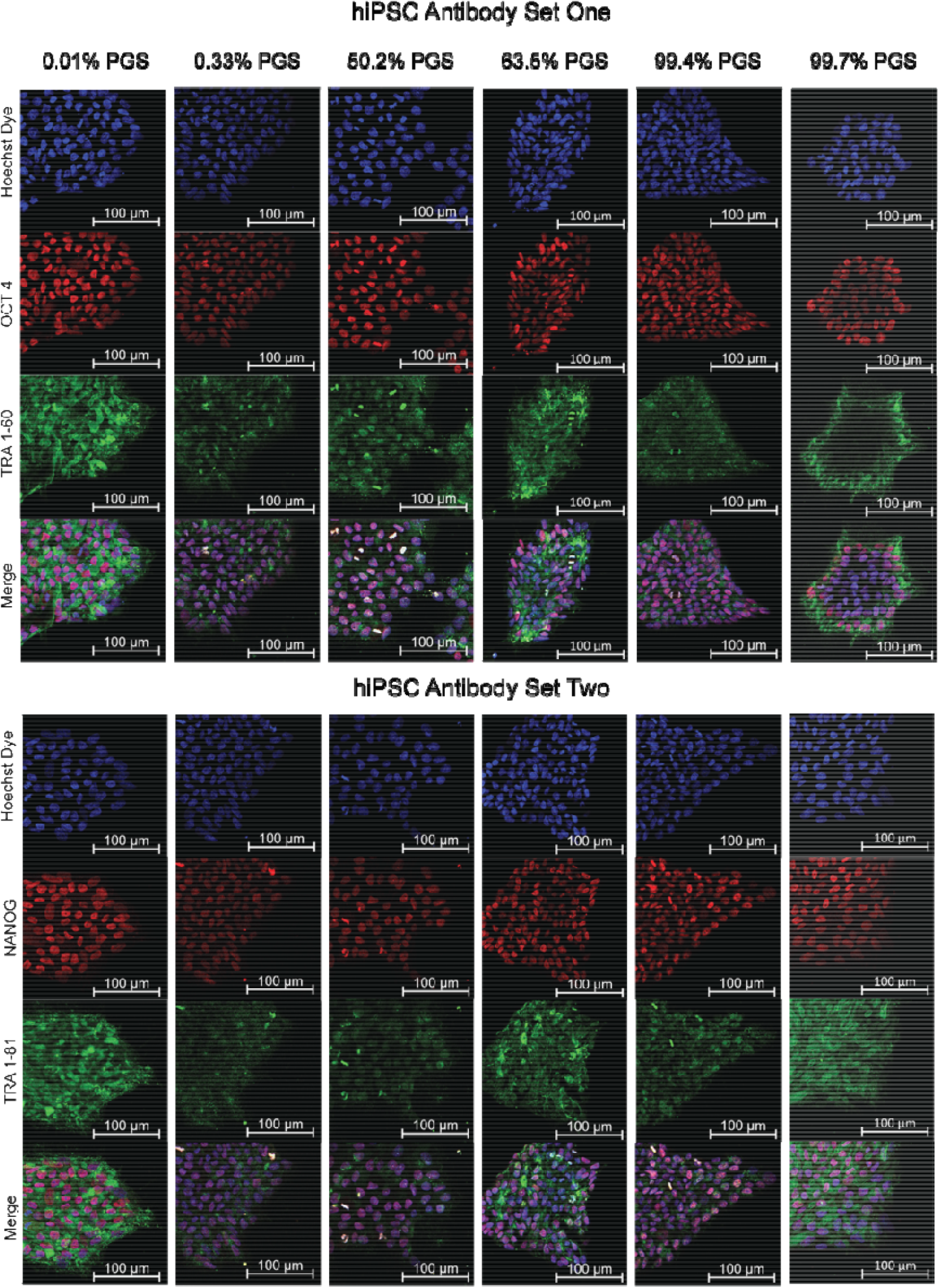
Immunofluorescence staining of six iPSC lines that display pluripotency makers: TRA 1- 60 antibody (green) is a common surface marker of stem cells, OCT 4 antibody (red) is a transcription factor highly expressed in undifferentiated stem cells, TRA 1-81 antibody (green) is a common surface marker of stem cells, and NANOG antibody (red) is transcription factor that maintains pluripotency. Both antibody sets have nuclei counterstained with Hoeschst dye (blue). The (0.01% and 033% PGS are low PGS iPSC lines, 50.2% and 63.5% PGS are medium PGS iPSC lines, 99.4% and 99.7% PGS are high PGS iPSC lines.)

### Culture of iPSC’s and cardiac induction

After each cell line was validated from the above processes, the cells were cultured with StemMACS(TM) iPS-Brew XF Basal medium (cat#130-107-086 Militenyi Biotec) in 6 well plates. When cells reached 65-70% confluency after approximately 4 days, the culture medium was aspirated and 1 mL per well of DPBS without Ca and Mg (Cat# 14190-144 Thermofisher Scientific) was added to wash the cells. The DPBS was then aspirated, and 1 mL per well of 0.5 mM EDTA (cat#46-034-CL Corning) was added and incubated for 8 min at room temperature. Meanwhile, 13 mL of fresh Brew medium with 13 μL 10 mM Rock inhibitor (cat#S1049 Selleckchem) was prepared in a 15 mL conical. After aspirating the EDTA from the well, 1 mL of the Brew+Rock inhibitor medium was added to the well and washed against the plate surface to break colonies into single cells. The cells were then equally plated into a 6-well plate (2 mL per well) and shaken back/forth and left/right 10 times each (repeat once for 20 times total). The plate was then placed in an incubator.

For differentiating iPSC’s into cardiomyocytes, we performed a series of media changes and additions of specific reagents at different time points. On day 0, cardiac induction was initiated by adding 3 mL of M1 (Table 1) with 6 µM of CHIR99021 (cat#S2924 Selleckchem) to the wells with cells at ∼55-65% confluency. On day 2, the medium was aspirated and 2 mL of M1 was added. On day 3, the medium was aspirated again, and 3 mL of M1 with 5 µM IWR-1 (cat#I0161 sigma) was added to the wells. On days 5, 7, and 9, the medium was changed by aspirating and adding 3 mL of M1. On day 10, metabolic selection was initiated by switching to M2 medium (Table 1). On days 12 and 14, the medium was changed by aspirating and adding 3 mL of M2. On day 15, the cells were redistributed by washing with DPBS, dissociated with 10x TrypLE Select (cat# A1217702 Gibco), and resuspended in DPBS and M2 media. The solution is then filtered with 100 μM sterile cell strainer (cat#22-363-549 Fisherbrand), and the cells were counted and centrifuged before being resuspended in 90% M2 medium and 10% FBS (Cat#A31606-01 Gibco). From day 16 to day 30, the medium was changed every other day by aspirating and adding 3 mL of M3 (Table 1) with 0.1 µM Triiodothyronine hormone (T3) (cat#T2877 sigma) and 1 µM Dexamethasone (cat#11015 Cayman). After day 30, only M3 medium was used to change the medium. The amount of T3 and Dexamethasone needed was calculated by adding the stock concentration to the M3 medium.

### Cardiomyocyte Confocal Microscopy

Fully differentiated Day 30 Cardiomyocytes were resuspended onto a cultrex stem cell qualified reduced growth factor basement membrane extract (cat# 3434-005-02 R&D systems Inc) coated 6-well plate with cover slips (cat#12-542-BP FisherBrand) in each well. For each 6-well added 2 mL of 4% PFA (cat#28906 Thermofisher) in PBS (cat#10010-023 Gibco) for 30 minutes at room temperature (RT). Following fixation, cells were washed three times with PBS to remove residual PFA. Next, cells were permeabilized with 0.2% Triton X-100 (cat#T8787 Sigma) in PBS for 20 minutes at RT. Cells were then blocked with 2 mL blocking solution made of 5% Normal Goat Serum (cat#PCN5000 Thermofisher) and 0.05% Triton X-100 in PBS for 2 hours at RT or 4°C on a shaker.

Primary antibodies against cardiomyocyte markers (Monoclonal Anti-α-Actinin antibody (Table 2), Myosin Light Chain 2/MLC-2V (Table 2) were added with 500 μL of blocking solution to the cells and incubated overnight at 4°C on a shaker. After incubation with primary antibodies, cells were washed three times with 0.05% Triton X-100/PBS for 5 minutes each. Secondary antibodies (Goat anti-Mouse IgG (H+L) Highly Cross-Absorbed Secondary Antibody Alexa Fluor 488 (Table 2), Goat anti-Rabbit IgG (H+L) Highly Cross-Absorbed Secondary Antibody Alexa Fluor 568 (Table 2) were then added with 500 μL of blocking solution and incubated for 4 hours at RT or overnight at 4°C on a shaker. After secondary antibody incubation, cells were washed three times with 0.05% Triton X-100/PBS for 5 minutes each. Next, 1 mL of Hoechst dye 2 ug/mL (cat#B1155 Sigma) in PBS were added to cells for nuclear counterstaining and incubated for 5 minutes at RT. After Hoechst dye incubation, cells were washed three times with PBS for 5 minutes each. Cells were then placed onto a desired glass slide (cat#12-550-433 Fisherbrand) with aqua hold 2 pen (Cat# 9804-02 Scientific Device) to help seal slide, and a cover slip is added with 50% glycerol (cat#17904 Sigma) in PBS. The edges of the cover slip were sealed with nail polish, and the slide was allowed to dry overnight at 4°C. Finished Stained hiPSC slides were then imaged on Zeiss LSM 710 confocal microscope. (Figure 3)

**Figure 3.**
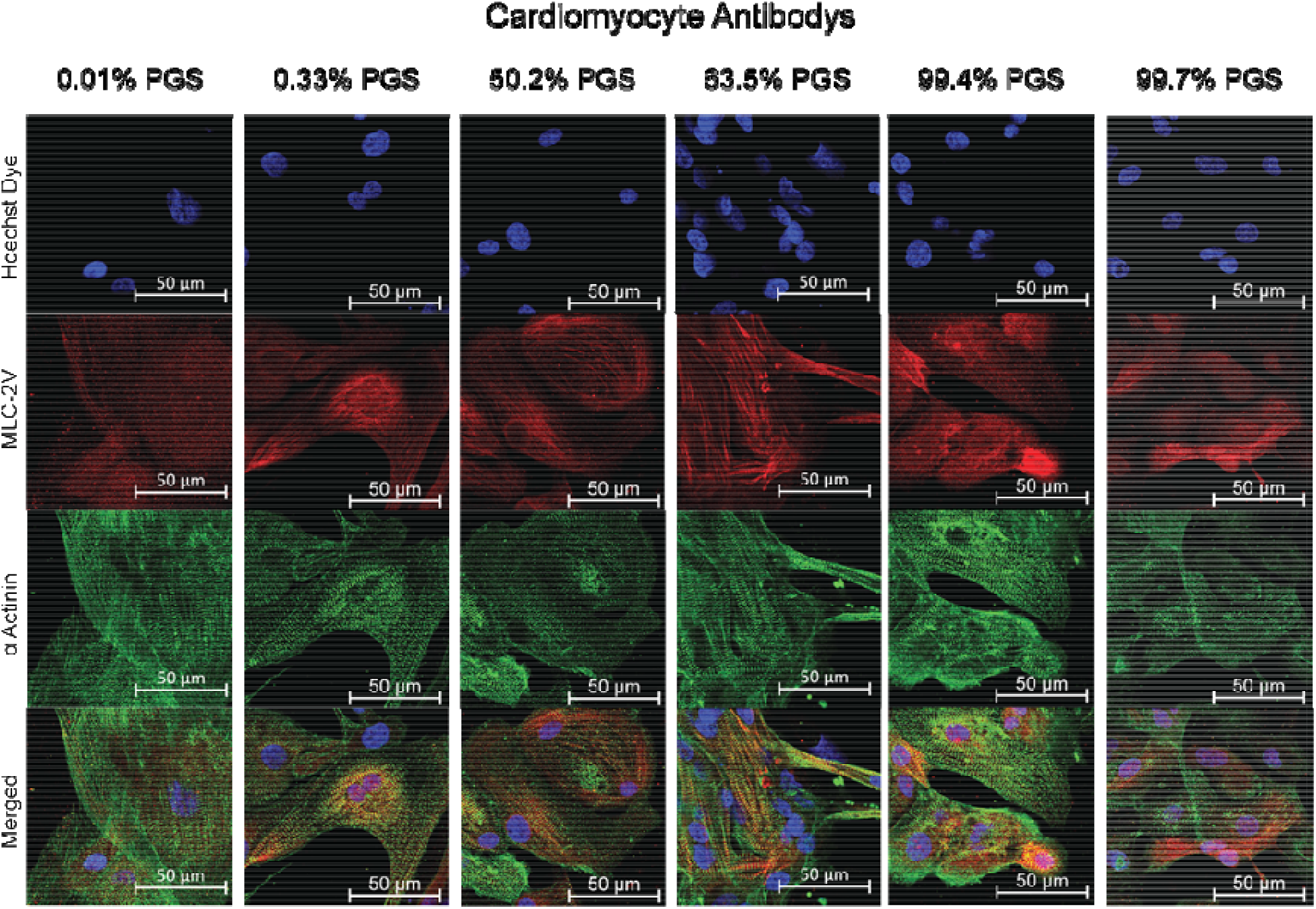
Immunofluorescence staining of six iPSC lines that displays cardiomyocyte markers: α- Actinin antibody (green) binds to cardiac actinin that shows striations throughout the cardiomyocyte and MLC-2V antibody (red) binds to myosin motor proteins that are specific to ventricular cardiomyocytes. Nuclei counterstained with Hoeschst dye (blue).

**Figure 4.**
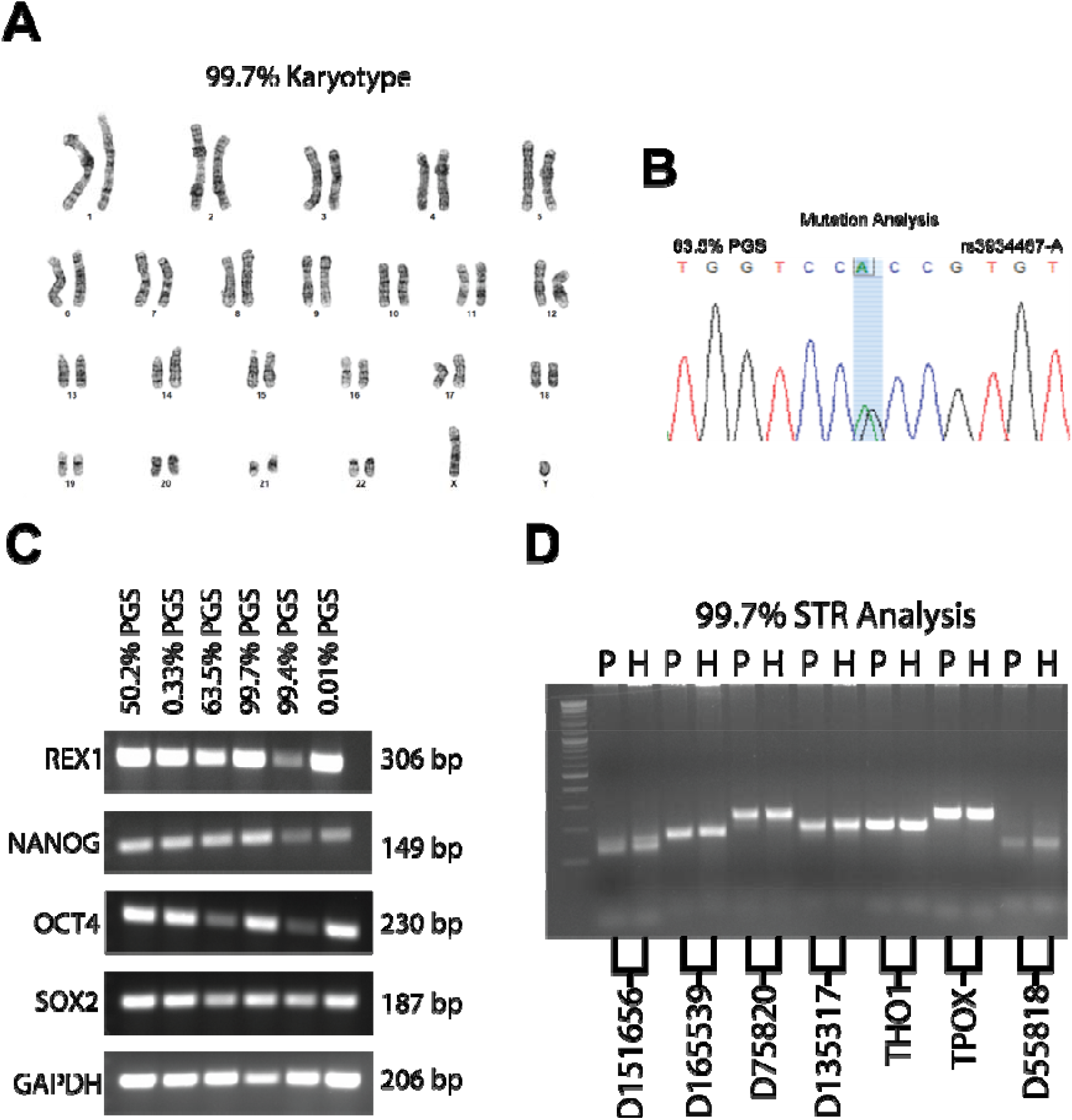
A. Karyotyping analysis of 99.7% PGS iPSC line displaying normal male karyotype. **B.** anger sequence results from a set of ten mutation loci, 63.5% PGS iPSC line shown as representative example of loci region for specific mutations to validate each reprogrammed hiPSC line. **C.** PCR of six iPSC lines to validate pluripotency marker expression of four commonly used markers (REX1, NANOG, OCT4, SOX2) and a house keeping gene (GAPDH). **D.** STR Analysis gel image of 99.7% PGS iPSC line containing seven commonly used loci bands amplified by PCR. P is DNA extracted from patient’s PBMCs for reprogramming, and H is DNA extracted from hiSPCs that have been reprogrammed from PBMCs.

