## Supplementary figures and images for "Generation of human induced pluripotent stem cell (hiPSC) lines from patients with extreme high and low polygenic scores for QT interval"

### Supplemental Material

**Chromosome images:**

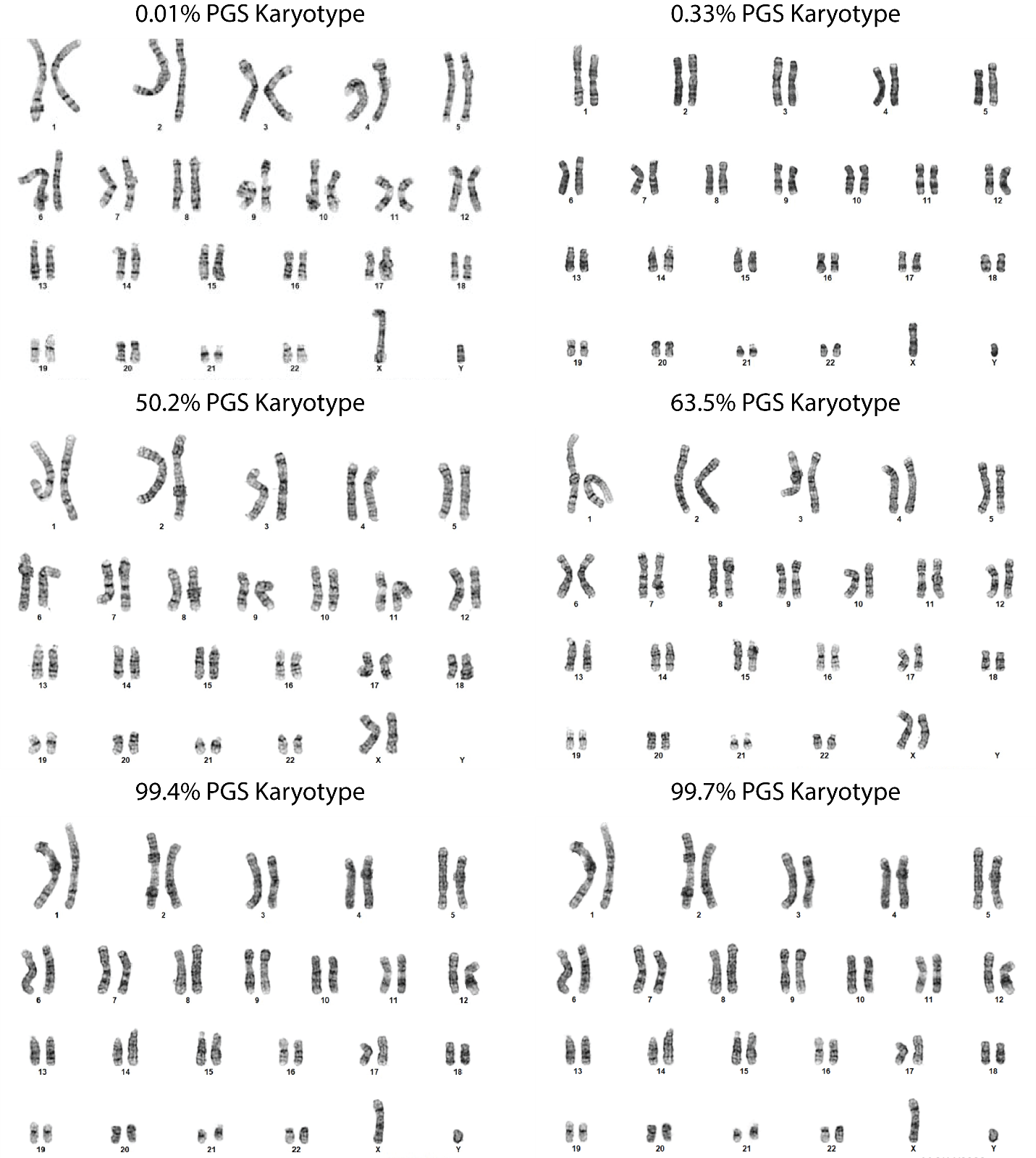

**Mutation Analysis images:**

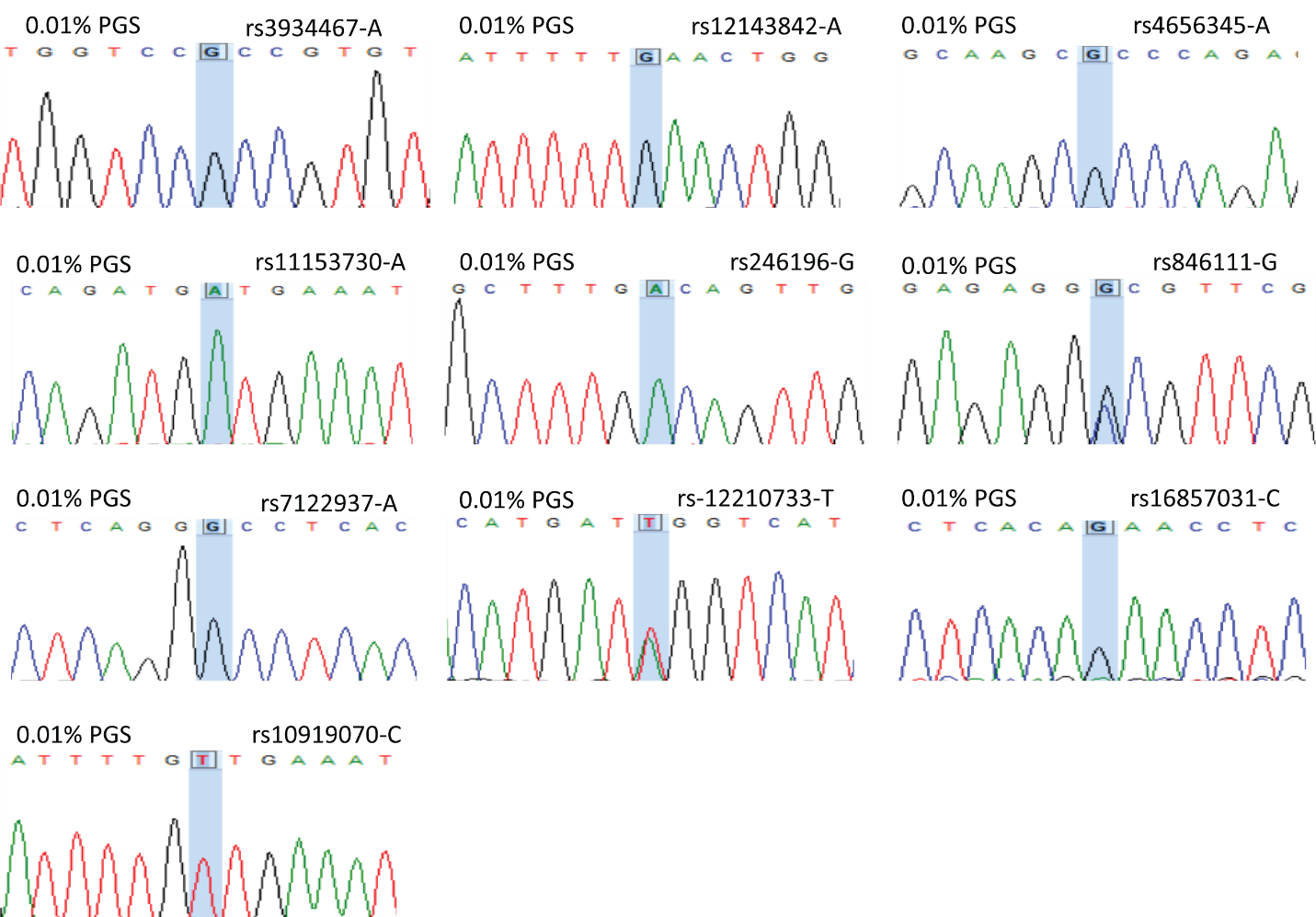

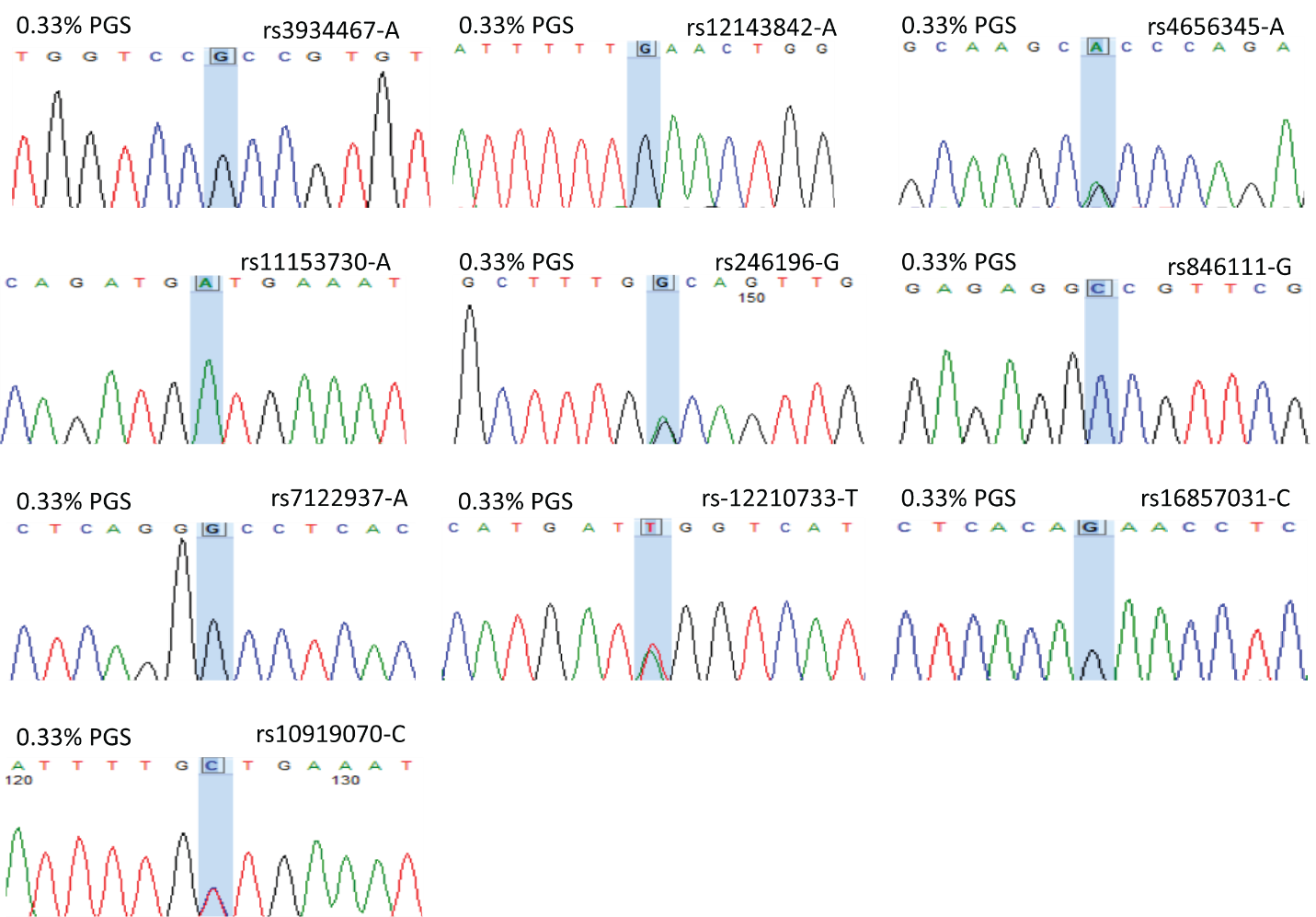

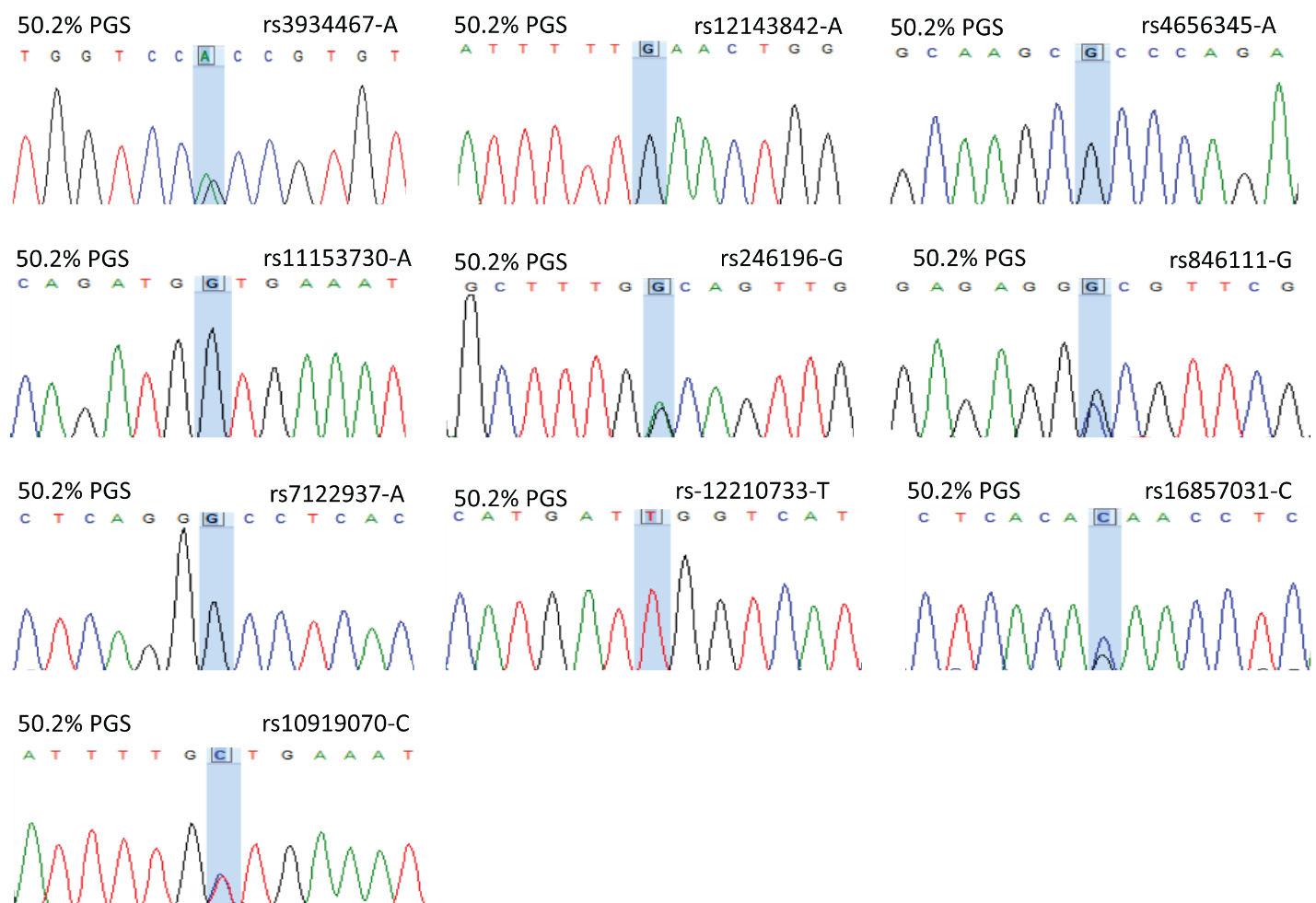

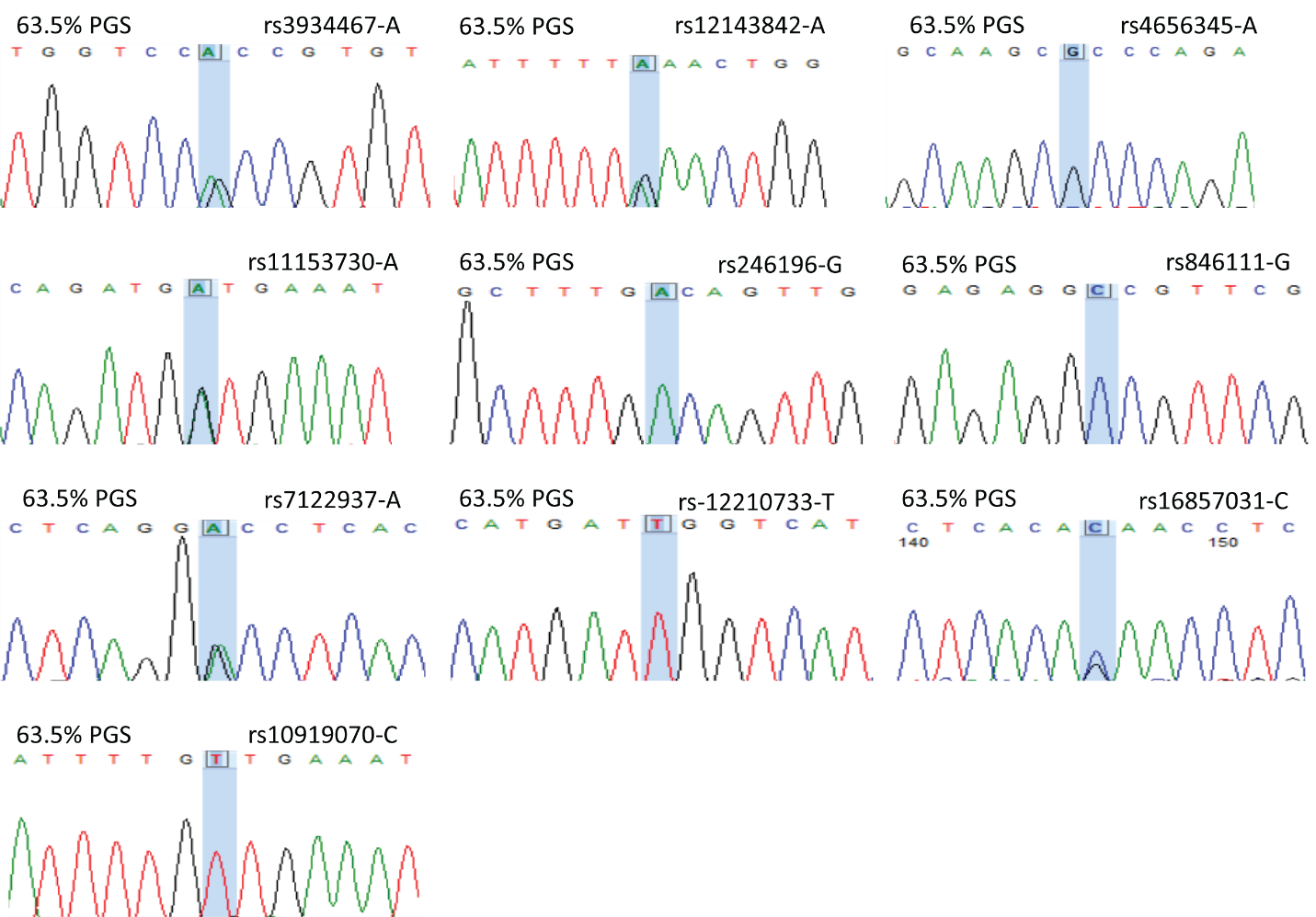

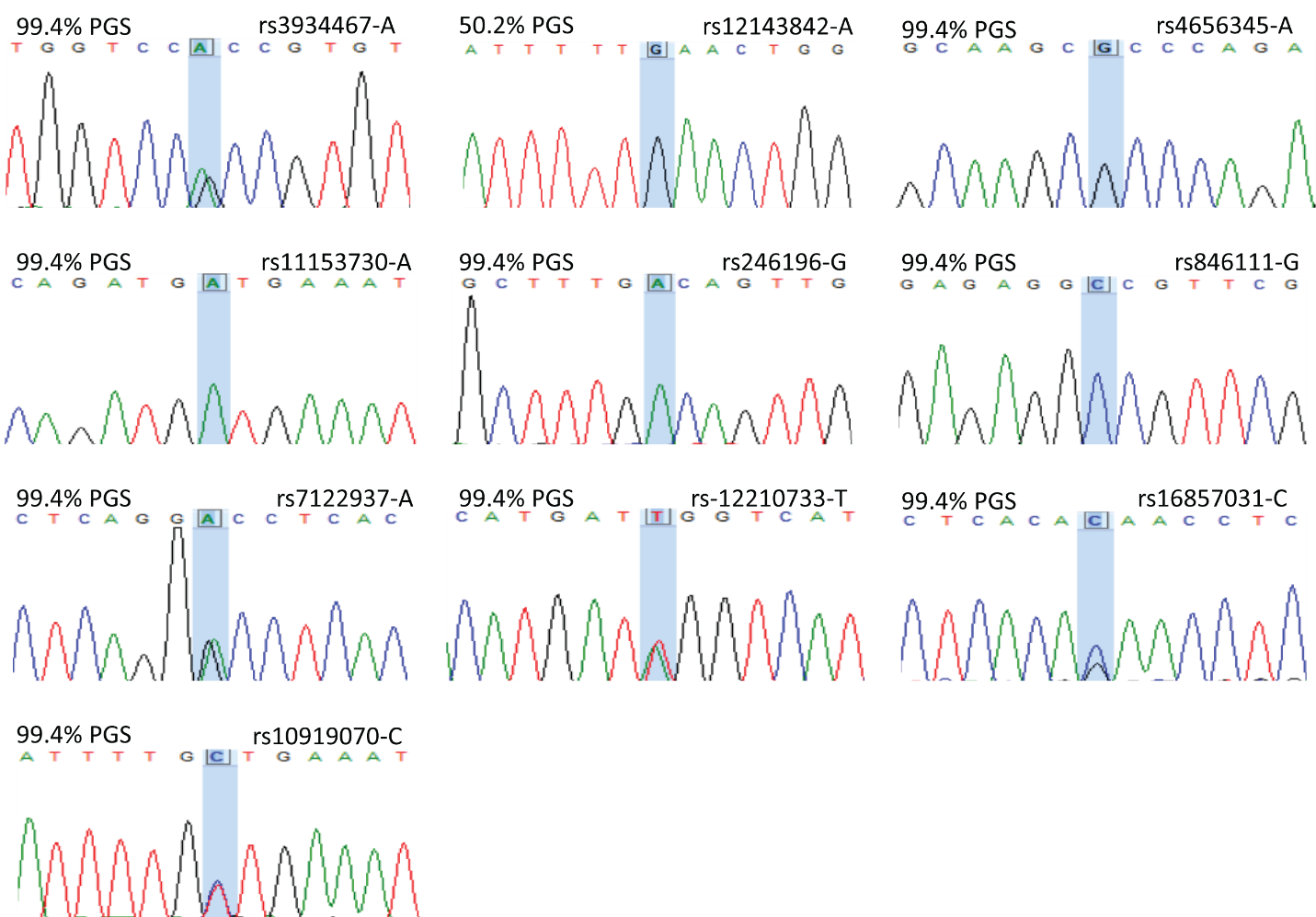

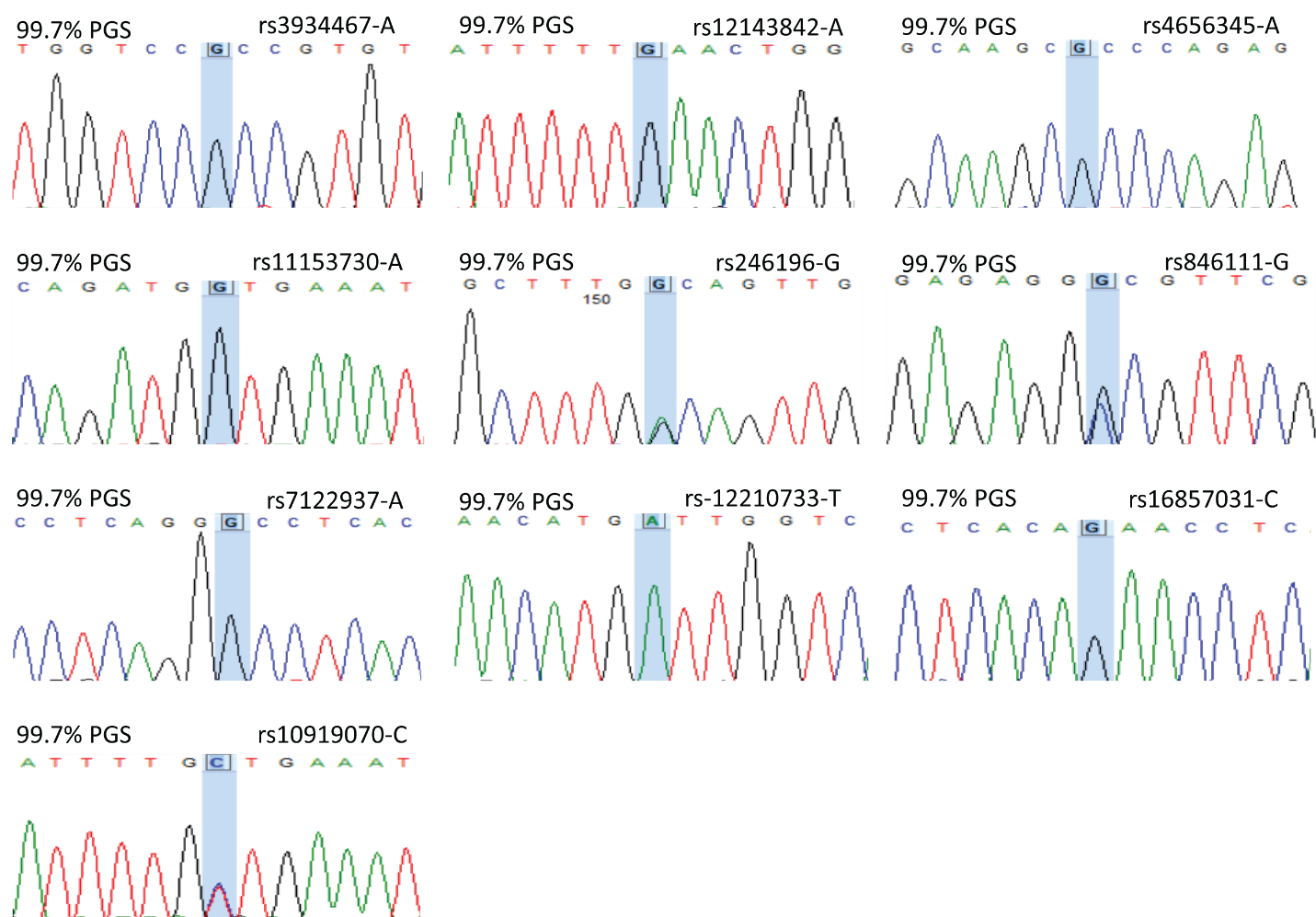

**STR Analysis images:**

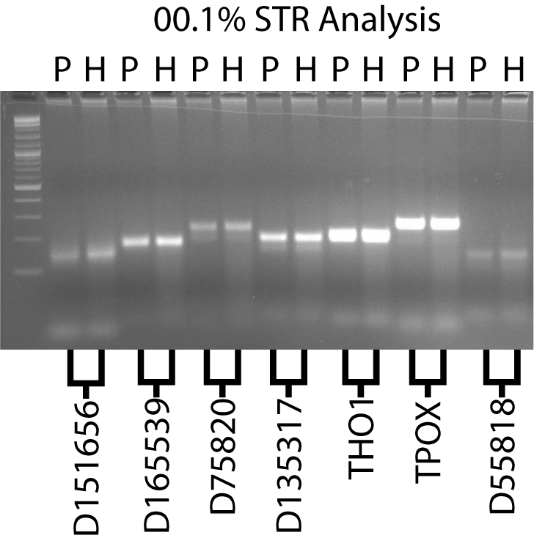

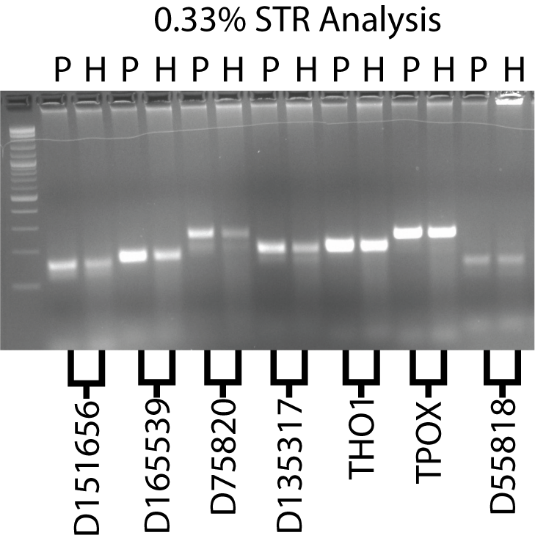

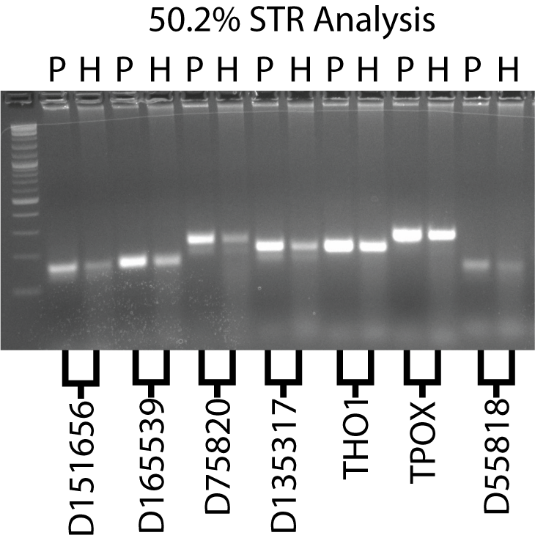

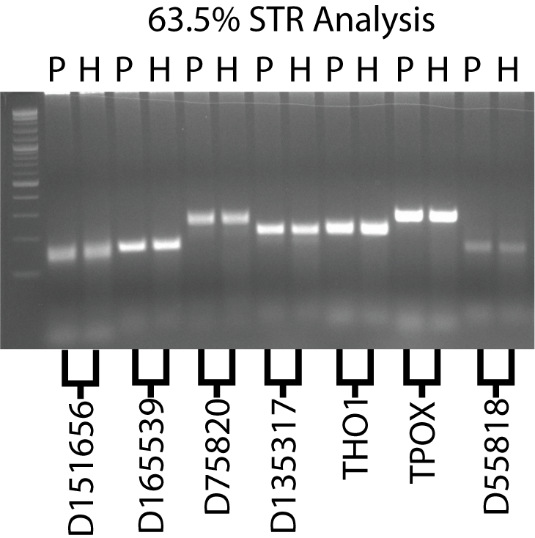

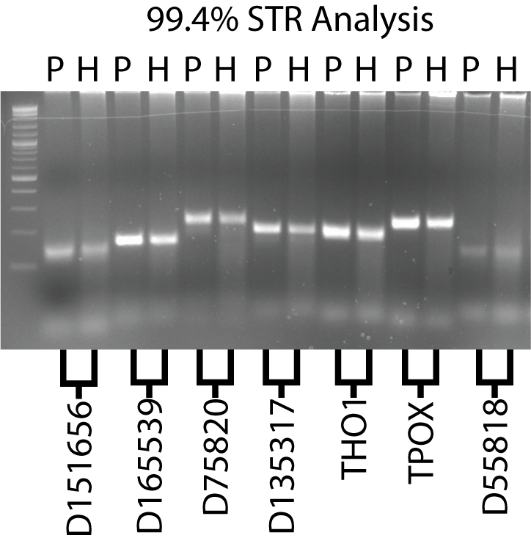

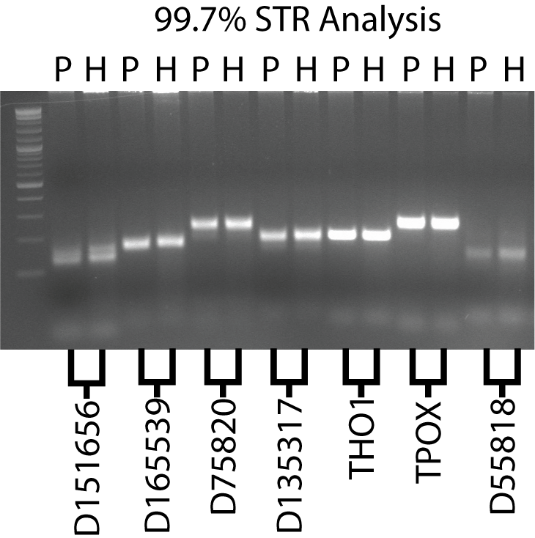
